# Unravelling the role of plant alpha-, beta-, and gamma- chemodiversity in shaping canopy insects of plants within the same species

**DOI:** 10.64898/2026.11.17.767245

**Authors:** Elikplim Aku Setordjie, Annika Neuhaus, Sunayana Chandra, Lina Ojeda-Prieto, Wolfgang Weisser, Robin Heinen

## Abstract

Plant intraspecific chemodiversity drives plant-herbivore and -pollinator relationships in plots, but whether it also impacts other members of the canopy arthropod community remains unclear. Furthermore, little is known of how different spatial components of plot intraspecific chemodiversity; chemical composition of individual plants within plots (i.e. α-chemodiversity); differences in chemical composition among plants within plots (i.e. β-chemodiversity); and cumulative chemical composition of all individuals in plots (i.e. γ-chemodiversity), impact aboveground community assemblage.

Using a chemodiversity field experiment containing 84 plots each planted with different combinations of Tansy (*Tanacetum vulgare* L., Asteraceae) chemotypes (i.e. a manipulated gradient of plot-level chemotype richness), we investigated the effect of plot-level chemodiversity and its components on the diversity and composition of canopy-associated arthropod communities within plots.

We found significant impacts of the presence of certain chemotypes on the effective number of common arthropod taxa (Hill q = 1) and the effective number of dominant arthropod taxa (Hill q = 2) within plots. Importantly, the observed pattern was independent of the actual terpenoid (α-)chemodiversity of these chemotypes. We also found that plot-level chemotype richness (β-chemodiversity) reduced the effective number of dominant arthropod taxa (Hill q = 2) within plots, driven by impacts on dominance of a nematoceran species associated with Tansy. Cumulative plot terpenoid richness (γ-chemodiversity) did not significantly impact arthropod communities. Instead, we observed that increased terpenoid abundance negatively influenced the effective number of common (Hill q = 1) and dominant arthropod taxa (Hill q = 2) within plots. Our results suggest that within plants of the same species, the chemotypic identity of plants and presence within plots and plot-level chemotype richness (β-chemodiversity) may impact arthropod communities by selecting for specific arthropod taxa making arthropod communities less even. Our results emphasize the nuanced impacts of chemodiversity at different levels and highlight taxon-specific arthropod responses.

## Introduction

Arthropod community assemblage is heavily influenced by the plant communities that host them (Ebeling et al., 2018; Scherber et al., 2010; Richards et al., 2015). These plant communities provide food, shelter and nesting sites for arthropods, but can also manipulate arthropod behaviour through the secondary metabolites they contain and emit (Bustos-Segura et al., 2017; Johnson & Gregory, 2006; Richards et al., 2015). Many studies have reported a positive relationship between inter- or intraspecific plant diversity and arthropod diversity (Bustos-Segura et al., 2017; Ebeling et al., 2018). Most studies with an intraspecific lens, however, focus on specific arthropod taxa or functional groups (e.g. herbivores), but much less is known about entire arthropod communities.

Intraspecific chemodiversity is an important component of plant biodiversity in ecosystems. This concept reflects how plants of the same species exhibit differences in the composition of particularly their specialized metabolites (Wetzel & Whitehead, 2020; Petrén et al., 2024; Ojeda-Prieto, Medina-van Berkum, et al., 2025). Such metabolomic differences influence plant ecological relationships and can be studied at different spatial or ecological scales, from plant individual, to community plots, to populations at landscape level (Bálint et al., 2016; Kleine & Müller, 2011; Neuhaus-Harr et al., 2025; Ojeda-Prieto, Moreno, et al., 2025; Setordjie et al., 2026; Ziaja & Müller, 2023). Many studies involve the use of intraspecific ‘chemotypes’: a plant or groups of plants belonging to the same species that are distinguishable by the profile of secondary metabolites within a chemical family in an organ (Müller et al., 2026). For instance, at plot-level, Sasidharan et al. (2024) observed higher pollinator visitations in heterogenous plots, containing different chemotypes, compared to homogenous plots with one chemotype. Similarly, in plots containing different combination of *Tanacetum vulgare*, Ojeda-Prieto, Moreno, et al., (2025), observed a positive relationship between plot-level chemotype richness and flower visiting arthropods, whilst herbivore abundance was negatively impacted. Chemotypic differences have also been found in the terpene concentration and diversity in *Monandra fistucola* plants in two different regions (Montana and Wisconsin) in the United States of America, resulting in associated differences in herbivore abundance and damage on plants varying in terpenoid diversity and their region of origin (Calixto et al., 2025).

Several hypotheses attempt to explain the existence of plant chemodiversity, many of them suggesting that plant chemodiversity is an evolutionary mechanism aimed at defence against abiotic and biotic stress (Wetzel & Whitehead, 2020). For example, the screening hypothesis posits that plants produce a large variety of phytochemicals as a means of increasing their odds of producing biologically active compounds that enhance their fitness, as the likelihood of producing active compounds is rare (Jones & Firn, 1991). In other words, organisms interacting with plants inadvertently select for plants with biosynthetic machineries that allow them to produce more diverse compounds since these plants are more likely to have active compounds (Jones & Firn, 1991; Wetzel & Whitehead, 2020). However, evidence for this hypothesis coming from natural environments is still scarce.

Plant intraspecific chemodiversity is often quantified using traditional biodiversity indices like species richness, Shannon diversity, and inverse Simpson diversity (sometimes calculated as Hill numbers) based on the richness (total number) and evenness (relative abundances/concentrations) of the different chemical compounds within a sampling unit, i.e. plant tissue, plant organ, individual plant, or plant community or population (Petrén et al., 2024; Wetzel & Whitehead, 2020). Similar to biodiversity, intraspecific chemodiversity can also be measured at different spatial scales using the α, β and γ diversity framework (Whittaker, 1972; Wetzel & Whitehead, 2020). In this case, α-chemodiversity measures the average chemical diversity within a single unit, which could be a tissue, organ, plant, plot, depending on the level of sampling. β-chemodiversity compares variations, dissimilarities, or turnover of specialized metabolites between different samples, and γ-chemodiversity pools chemical diversity of different samples across their respective group, for instance across plant communities, populations or regions (Petrén et al., 2024; Schneider & Beckman, 2025; Wetzel & Whitehead, 2020).

Although research on intraspecific plant chemodiversity and its effects on ecological relationships has gained traction in recent years (Calixto et al., 2025; Ojeda-Prieto, Moreno, et al., 2025; Sasidharan et al., 2024; Ziaja & Müller, 2023; Bustos-Segura et al., 2017; Glassmire et al., 2020; Kleine & Müller, 2011), the lack of a uniform, standardised method of estimation means inferences drawn across studies can be challenging (Petrén et al., 2024; Wetzel & Whitehead, 2020). Additionally, with most studies of chemodiversity focusing on the ecological consequences of chemodiversity on closely associated organisms like herbivores and pollinators, the numerous more transient arthropod groups found in plant canopies are often neglected.

Here, we make use of a field experiment containing 84 plots, each planted with six Tansy (*Tanacetum vulgare*) plants, an Asteraceae plant characterised by its high intraspecific mono- and sesquiterpenoid diversity (Rahimova et al., 2024). We varied chemotype proportions within plots to achieve chemotype richness levels of 1, 2, 3 and 6, resulting in a clear increase in predicted plot-level leaf terpenoid chemodiversity (Ojeda Prieto, Medina van Berkum, et al., 2025). Using a vacuum suction sampler, we sampled the aboveground arthropod community in each plot. We used known terpenoid profiles for plants in the field experiment as proxies to examine the effect of α-chemodiversity (presence or absence of different Tansy chemotypes within plots, ranking chemotypes by their average terpenoid diversity); β-chemodiversity (dissimilarities in leaf terpenoid and plant chemotype composition between plots); and γ-chemodiversity (cumulative leaf terpenoid diversity of all plants within plots) on aboveground arthropod assemblage (Figure 1).

**Figure 1:**
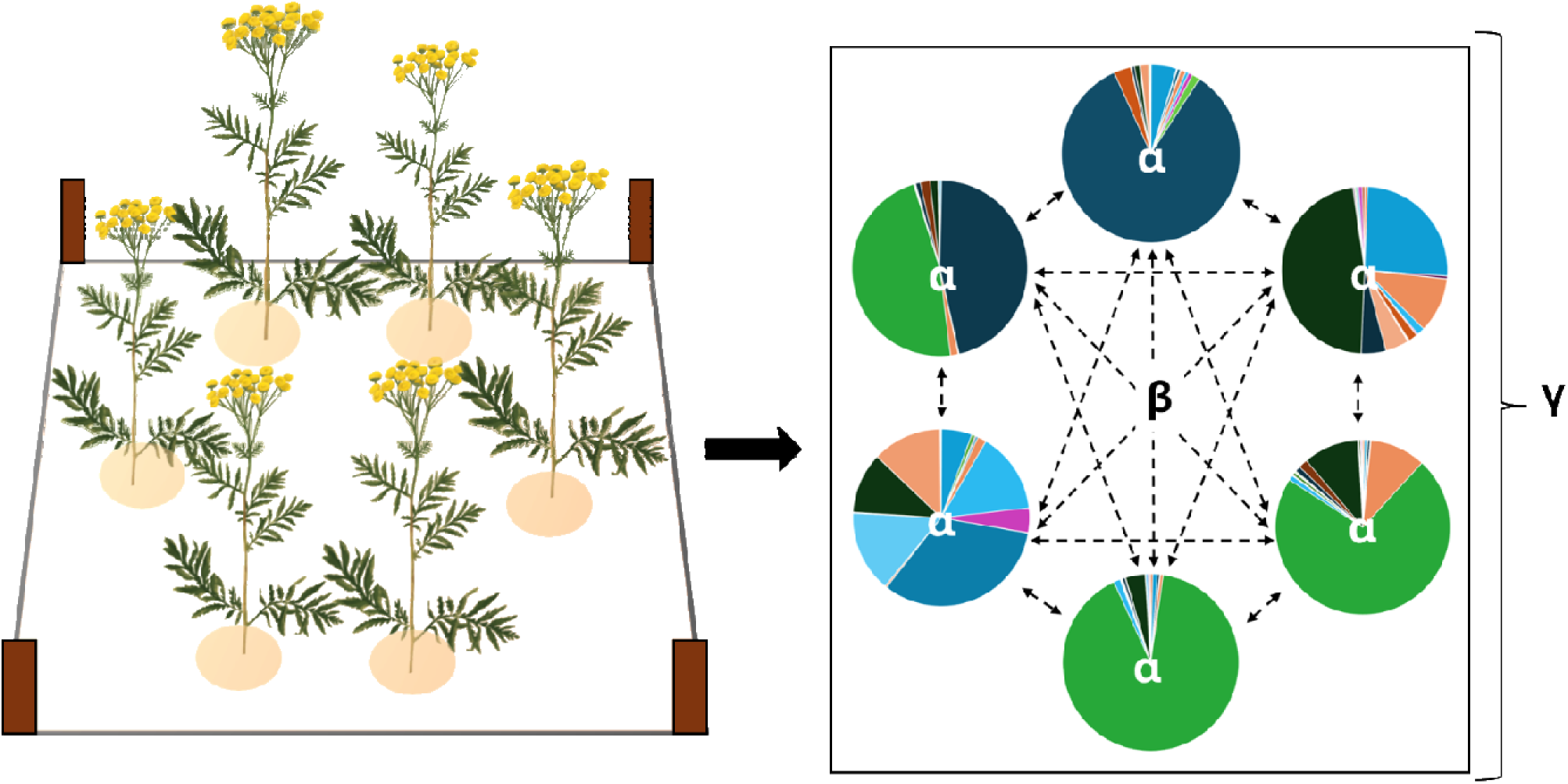
Graphical representation of α, β and γ-chemodiversity within plots, where circles (pie charts) represent individual Tansy plants, partitions within circles represent α-chemodiversity i.e. the different leaf terpenoids and their abundances in individual *Tanacetum vulgare* (Tansy) plants, dashed arrows show β-chemodiversity i.e. differences/turnover in leaf terpenoid composition between plants within plots and all circles present within box represent γ-chemodiversity i.e. total leaf terpenoid diversity within plot.

We specifically tested the following hypotheses:

**H1:** The presence of specific Tansy chemotypes within plots alters associated aboveground arthropod communities (Bálint et al., 2016; Neuhaus-Harr et al., 2024). The presence of chemotypes with more variable terpenoid profiles (i.e. higher α-chemodiversity) will result in more diverse associated arthropod communities within plots whilst plots containing chemotypes with less variable terpenoid (i.e. lower α-chemodiversity) profiles attract less diverse arthropod communities.

**H2:** Plots containing more similar terpenoid profiles (β-chemodiversity) will attract more similar arthropod communities. If terpenoids evolved as means of defence against herbivores, then the presence of a particular terpenoid should serve as a cue for its co-evolved arthropod group. Therefore, plant communities (in this case plots) with similar terpenoid profiles are expected to have similar associated arthropod communities and plots having plants with more dissimilar terpenoid profiles should have more dissimilar arthropod taxa (Wetzel & Whitehead, 2020).

**H3:** Higher γ-chemodiversity at plot-level leads to higher arthropod diversity. Having a higher number of chemicals i.e. terpenoids, within a plot increases the chances of the plot attracting a broader number of arthropod species, which may be adapted to varying chemical components of the plot as posited by the screening hypothesis (Wetzel & Whitehead, 2020).

## Materials and Methods

### Experimental design

To describe tansy plant chemotypes their leaf terpenoid concentrations were subjected to cluster analysis and named after their dominant terpenoids (that is, “Athu-Bthu”, “Bthu-low”, “Bthu-high”, “Mixed-high”, “Mixed-low” and “Chrys-acet”) (as in Neuhaus-Harr et al., 2024; Ojeda-Prieto et al., 2025). For each chemotype, three unique daughters were selected and clonally propagated.

The experimental field was set-up in 2021 next to the Jena grassland biodiversity Experiment in Jena, Germany (Weisser et al., 2017) and has been maintained ever since. It consists of 84 plots (1m × 1m, separated by 70 cm wide foot paths), each planted with six plants. Plants were chosen from 6 Tansy chemotypes to attain chemotype richness levels of 1, 2, 3 or 6 within a plot (Fig. 1). Plots were distributed within 6 replicate blocks containing 14 plots each. Chemotype richness levels were replicated 12 times for levels 1 and 6 and 30 times for levels 2 and 3 across the field (Fig. S1). In the choice of specific daughters for each chemotype, we set-up different unique plot-chemotype combinations in the experiment (i.e. each of the resulting 18 daughters were present 27-31 times across the field. The unique plot compositions are presented in Ojeda-Prieto et al., (2025).

### Arthropod community sampling

The aboveground arthropod community within each plot was sampled on the 13^th^ of August and the 14^th^ of August 2024 (3 blocks sampled per day) at average daily temperatures between 25°C and 26°C (Deutscher Wetterdienst, 2024) obtained using the ‘*rdwd’* package in R (Boessenkool, 2017). Plants within each plot were carefully tied upright with a rope the day before sampling to allow for minimal disturbance while installing the sampling cages and to allow sufficient time for arthropod communities to recolonise plants. On the day of sampling a 1 m × 1 m × 1 m netted aluminium cage frame was placed over each plot. Then, using a modified commercial vacuum cleaner (Kärcher WD3, Kärcher GmBH, Winnenden, Germany) with nozzle fitted with a perforated plastic bag and secured with a rubber tie, the aboveground arthropod community was sampled for four minutes per plot (for optimal coverage, the same sampler suctioned from opposite ends of the cage for two minutes each, Fig. S2). The arthropod samples collected were transferred into 70% ethanol and transported to the laboratory for sorting into order and suborder. Lower taxonomic resolutions were obtained for the orders with highest abundance; Diptera: Brachycera, Nematocera (Cecidomyiidae), Hemiptera: Heteroptera, Auchenorrhyncha, Psyllidae, Aphididae, Hymenoptera: Symphyta, Apocrita, Formicidae.

*Arthropod abundance* per plot was determined by summing up all individual arthropods sampled within the plot, whilst *arthropod diversity* metrics were determined using Hill numbers, i.e. q = 0 (*Arthropod taxon richness*), q = 1 (Shannon diversity, i.e. *Effective number of common taxa*) and q = 2 (Inverse Simpson diversity, i.e. *Effective number of dominant taxa*) calculated using the identity and numbers of arthropod taxa found within each plot (Chao et al., 2021).

### Terpenoid diversity metrics

Plot chemodiversity metrics were calculated as Hill numbers, i.e. q = 0 (*Terpenoid richnes*s), q = 1 (Terpenoid Shannon diversity, i.e. *Effective number of common terpenoids*) and q = 2 (Terpenoid inverse Simpson diversity, i.e. *Effective number of dominant terpenoids*) based on the cumulative leaf terpenoid profiles of each plant present within the plot. The leaf terpenoid concentrations (nmol/g) were derived from available leaf terpenoid profiles from the 18 daughters used to establish the field in 2021 (Ojeda-Prieto, Medina-van Berkum, et al., 2025), which are used as proxies for terpenoid representation within the plots. Leaf samples from these plant lineages have been chemotyped multiple times since from cuttings originating from the same original mothers, and terpenoid profiles have been shown to remain consistent across different experimental years (Newrzella et al., 2026). *Plot terpenoid abundance* was estimated by adding up the concentrations of terpenoids present within a plot based on the available leaf terpenoid concentrations standardized per unit leaf biomass.

### Plant morphological traits

Larger plants provide more resources and space to arthropods, which can also increase the abundance and diversity of associated arthropods independent of chemotype. To account for the influence of plant size on arthropod communities, we measured two morphological traits, plant height and average stem numbers, to be used as covariates in the analyses. The number of stems per plant was recorded once in July 2024 whilst the height of the longest stem per plant was recorded once in August of 2024. Both traits were averaged over the six plants per plot and referred to as *average number of stems* within plots and *average plant height* within plot respectively.

### Statistical analysis

All statistical analyses were performed using R version 4.5.0 (R Core Team, 2025). To visualise plant terpenoid variation and distribution across different plot chemotype richness levels, a principal component analysis was performed on the log-transformed terpenoid concentrations of plants within each plot using the ‘*vegan*’ package (Oksanen et al., 2025) (Fig. S3).

Plot terpenoid diversity metrics were estimated using the *‘chemodiv’* package (Petrén et al., 2023), whilst arthropod diversity within plots was determined by calculating hill numbers (q = 0,1,2) using the ‘*hillR*’ package (Li, 2018).

To test for **H1** (α*-chemodiversity effects on arthropod community*), we evaluated the (fixed) effect of the presence of individual Tansy chemotypes on arthropod taxa diversity, i.e. q = 0,1,2, and arthropod abundance using linear mixed effects models via the ‘*lme4*’ package (Bates et al., 2015). Block was included as a random effect to correct for spatial variability within the field, as well as the sampling design which incorporated three blocks per sampling day. To qualitatively assess the effect of α-chemodiversity, we ordered the results according to the mean α-chemodiversity of each Tansy chemotype.

To test H**2** (β*-chemodiversity effects on arthropod community*) we used three approaches. We first compared pairwise dissimilarity matrices in terpenoid and arthropod composition per plot.

For this, we calculated pairwise similarity between plots in both terpenoid composition, i.e. leaf terpenoids of plants present within plots and their concentrations, and arthropod community composition, i.e. arthropod taxa present within plots and their abundances, using Bray-Curtis dissimilarity within the *‘vegan’* package, followed by a Mantel test, ran at 9999 permutations to determine the relationship between the two dissimilarity matrices.

As a second approach, we tested the effect of the similarity of the chemical composition of plants within the plots on arthropod taxa diversity, q = 0,1,2, and arthropod abundance. For this, we calculated the pairwise dissimilarity between individual plants within each plot in terpenoid composition (leaf terpenoid identity and abundance) using the *‘vegan’* package. *Average plot dissimilarity* was estimated by adding up all the pairwise dissimilarities within each plot and dividing by the total number of plants present within the plot (i.e. 6). The (fixed) effect of Average plot dissimilarity on arthropod taxa diversity, q = 0,1,2, and arthropod abundance was then determined using linear mixed models via the ‘*lme4*’ package (Bates et al., 2015). Block was included as a random effect in mixed models to correct for spatial variability within field.

Thirdly, we assessed the (fixed) effect of plot-level chemotype richness on arthropod taxa diversity, q = 0,1,2, and arthropod abundance using linear mixed models with the same random effect structure described above.

To evaluate **H3** (γ*-chemodiversity effects on arthropod community*), the (fixed) effect of plot terpenoid diversity, q = 0,1,2, on arthropod taxa diversity and arthropod abundance was assessed using linear mixed models. Block was included as a random effect to account for the spatial structure of our experimental design, as well as the sampling design which incorporated three blocks per sampling day. We also evaluated the (fixed) effect of plot terpenoid abundance on arthropod taxa diversity and arthropod abundance using the same random effect structure described above.

General structure of all models:

~~~
lmer(Arthropod community metric ∼ Average number of stems within plots +
Average plant height within plots + Plot chemodiversity metric +(1|Block))
~~~

Model specifications are included in the supplementary information of the manuscript. Data was organised using ‘*tidyverse*’ (Wickham et al., 2019). All graphics were created using ‘*ggplot2’* (Wickham, 2016) and ‘*patchwork*’ (Pedersen, 2019). All *p*-values were calculated using the Anova II function in the ‘*car’* package (Fox et al., 2019).

## Results

Overall, we documented a total of 8624 arthropods within our field from 20 (sub)orders. The dominant (sub) orders observed in our plant canopies were Apocrita, Nematocera and Heteroptera with 3368, 2883 and 1159 individuals, respectively. Larvae of holometabolous insects were excluded from the analysis due to rarity of occurrence (2 individuals) and the difficulty to place them accurately in an order or suborder (Fig. S5).

### Influence of tansy chemotype and α-chemodiversity on arthropod community diversity and abundance

The presence of all Tansy chemotypes did not significantly affect arthropod taxon richness within plots (Fig. 2a, Table S1).

**Figure 2:**
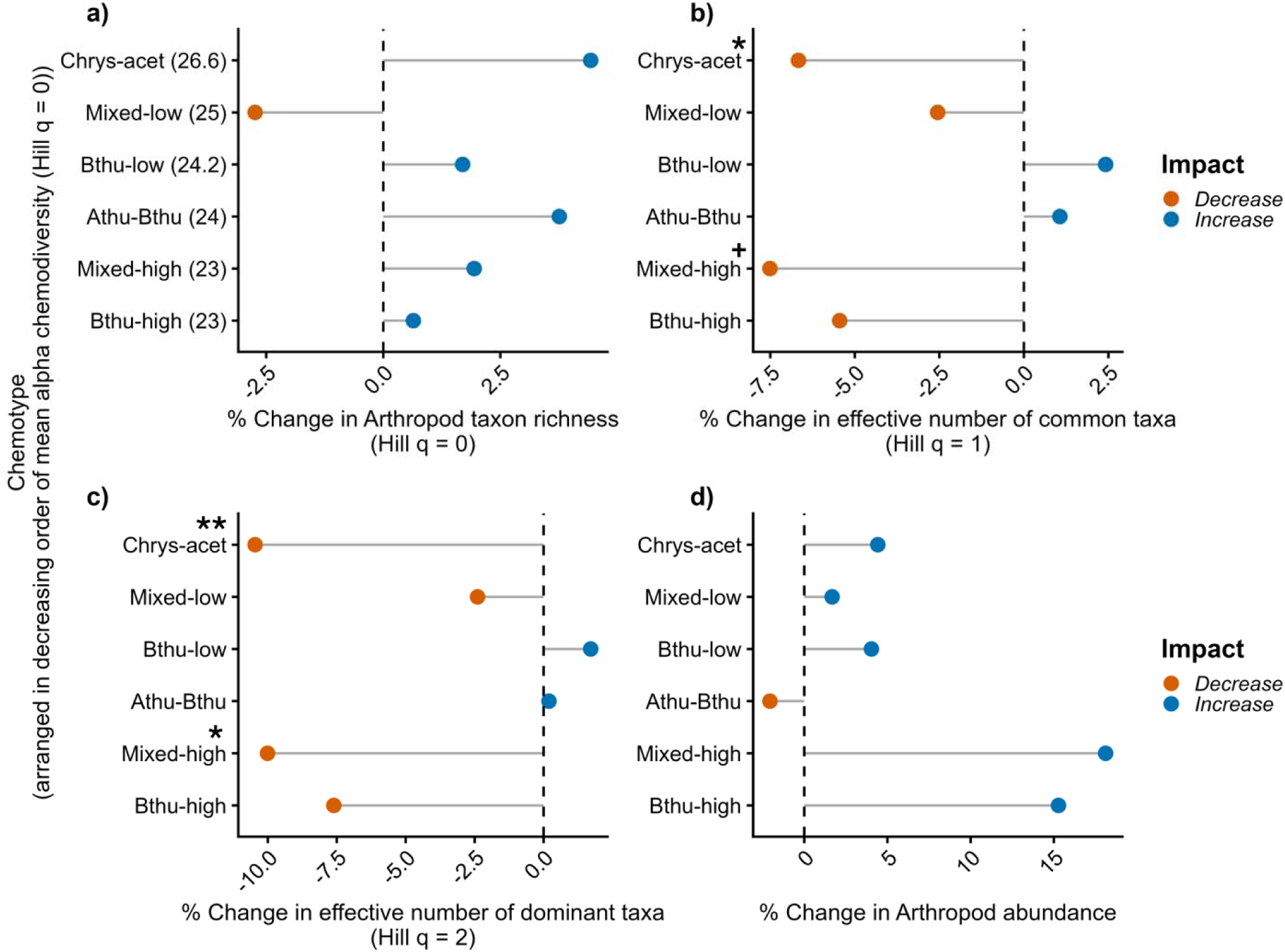
Effect of the presence of Tansy chemotypes on percentage change in (a) Arthropod taxon richness (Hill q = 0), (b) Effective number of common arthropod taxa (Hill q = 1), (c) Effective number of dominant arthropod taxa (Hill q = 2) and (d) Arthropod abundance. Numbers between parentheses after the chemotypes in panel (a) represent the mean alpha-chemodiversity (Hill q = 0). Chemotypes are arranged in decreasing order from top to bottom. Significance levels are presented as: + = 0.10 > *p* > 0.05, * *p* <0.05, ** *p* <0.01, and *** *p* <0.001 on top of chemotype names.

The presence of most Tansy chemotypes did not significantly influence the effective number of common arthropod taxa recorded in plots except for the Chrys-acet (*X*² = 4.2, p = 0.041) and Mixed-high (*X*² = 3.6, p = 0.056) chemotypes, whose presence reduced the effective number of common taxa observed in plots by 6.7% and 7.5% respectively (Fig. 2b, Table S1).

Presence of the Chrys-acet (*X*² = 7.0, p = 0.008) and Mixed-high (*X*² = 4.8, p = 0.029) chemotypes within plots also led to a decline in the effective number of dominant taxa observed within plots by 10.5 % and 10.0 % respectively (Fig. 2c, Table S1).

The average number of stems and the average plant height within plots did not significantly influence arthropod diversity (i.e. Hill q = 0,1,2), in the presence of all Tansy chemotypes (Table S1).

With respect to total arthropod abundance, plots with higher average plant height had increased arthropod abundance regardless of which Tansy chemotypes were present (Table S1). The presence of a particular Tansy chemotype and the average number of stems within plots did not significantly influence arthropod abundance within plots (Fig. 2d, Table S1).

Ranking of chemotypes by their respective mean alpha diversity revealed no obvious trend between the mean α-chemodiversity of respective chemotypes and the diversity and abundance of arthropod taxa (Fig. 2).

### Influence of plot β-chemodiversity on arthropod community diversity and abundance

#### Relationship between plot terpenoid composition and plot arthropod composition

Dissimilarities between plot terpenoid composition (that is, the leaf terpenoid identity and concentration of plants within plots) were not significantly correlated with dissimilarities in arthropod taxa (arthropod taxa present within plots and their abundances) (Mantel r = 0.008, p = 0.42) (Fig. S4).

#### Relationship between terpenoid dissimilarity between plants in plots and arthropod diversity and abundance

Average plot terpenoid dissimilarity, average number of stems and average plant height in a plot did not significantly affect arthropod taxa diversity (q = 0,1,2) (Table S2).

In contrast, arthropod abundance was positively influenced by average plant height within plots (*X*² = 5.2, p = 0.023), but not by plot terpenoid dissimilarity (*X*² = 0.7, p = 0.391) and average number of stems (*X*² = 0.0, p = 0.978) (Table S2, Fig. S6).

#### Plot-level chemotype richness effect on arthropod diversity and abundance

Arthropod taxon richness (Hill q = 0) and the effective number of common taxa (Hill q = 1) within plots were not significantly affected by plot-level chemotype richness, average number of stems and average plant height (Fig. 3a and 3b, Table S3).

**Figure 3:**
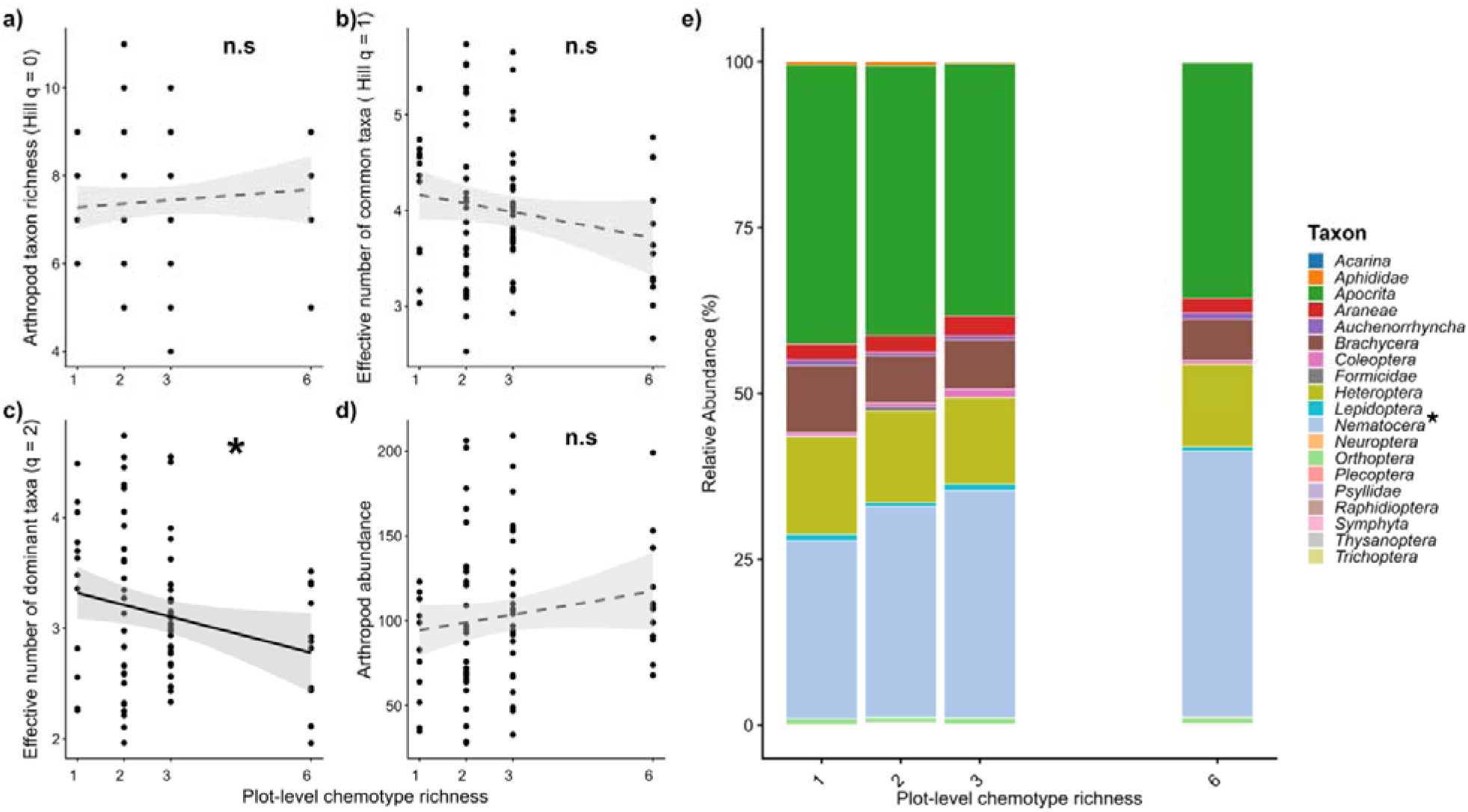
Effect of plot-level chemotype richness on (a) Arthropod taxon richness (Hill q = 0), (b) Effective number of common taxa (Hill q = 1), Effective number of dominant taxa (Hill q = 2) and (d) Arthropod abundance. e) Distribution of arthropod taxa across different Tansy plot chemotype richness levels. Significance levels are presented as: n.s = not significant, + = 0.10 > *p* > 0.05, * *p* <0.05, ** *p* <0.01, and *** *p* <0.001.

Similarly, the average number of stems (*X*² = 0.3, p = 0.556) and the average plant height (*X*² = 1.3, p = 0.252) had no significant impact on the effective number of dominant taxa (Hill q = 2) found within plots. Plot-level chemotype richness (*X*² = 4.0, p = 0.045) however, led to a decrease in the effective number of dominant taxa observed in plots (Hill q = 2, Fig. 3c and e, Table S3).

Plots with higher average plant height (*X*² = 4.1, p = 0.043) had a higher arthropod abundance. Neither plot-level chemotype richness (*X*² = 1.7, p = 0.192) nor average number of stems within plots (*X*² = 0.0, p = 0.885) influenced arthropod abundance (Fig. 3d, Table S3).

### Influence of plot γ-chemodiversity on arthropod community diversity and abundance

#### Plot-level terpenoid diversity metrics effects on arthropod diversity and abundance

Terpenoid diversity (Hill q = 0,1,2), terpenoid abundance, average number of stems and average plant height within plots had no significant effect on arthropod taxon richness (Hill q = 0) in plots (Fig. 4a, S7a, S8a, S9a & Table S4).

**Figure 4:**
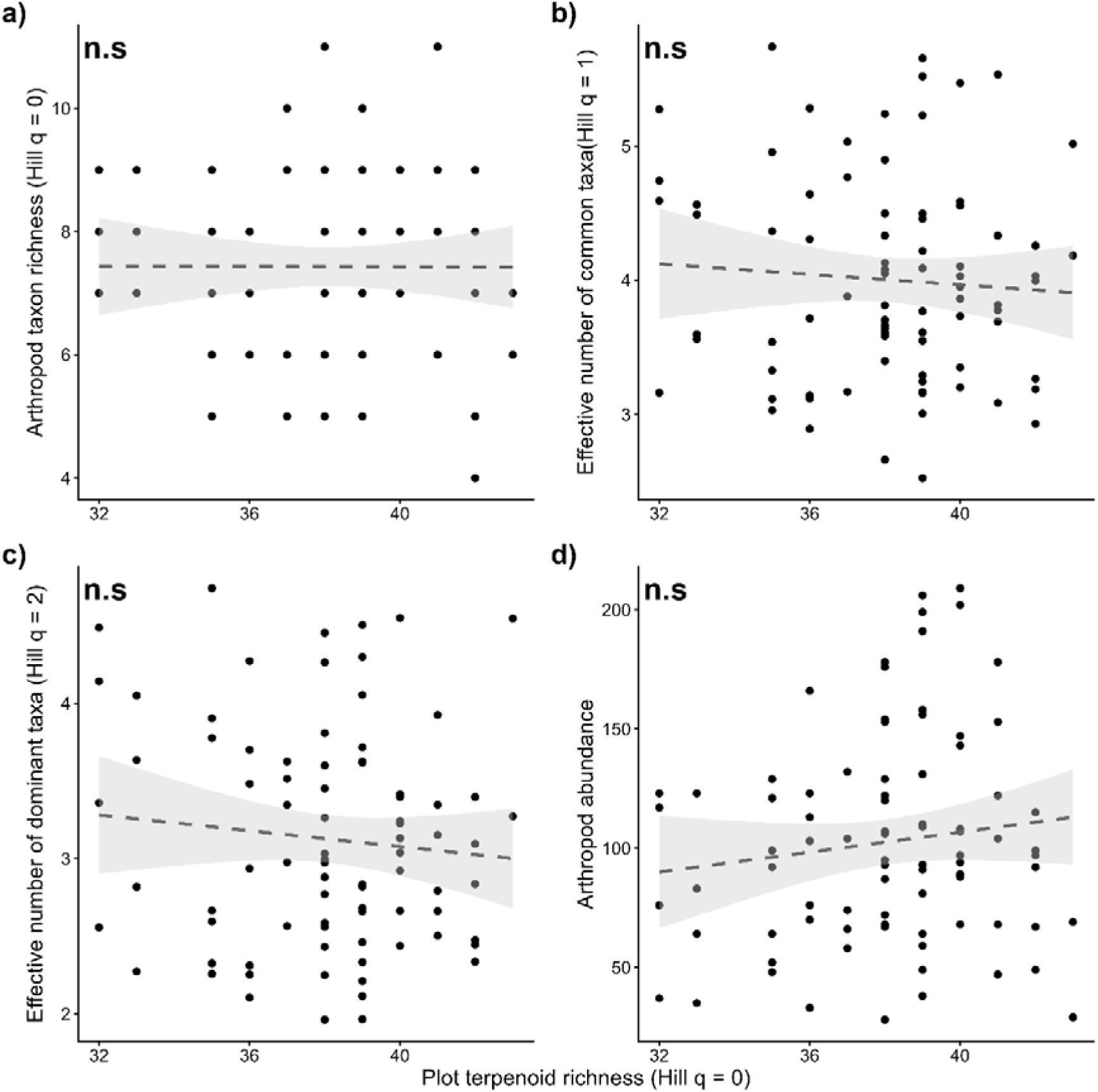
Effect of plot terpenoid richness (Hill q = 0) on (a) Arthropod taxon richness (Hill q = 0), (b) Effective number of common taxa (Hill q = 1), (c) Effective number of dominant taxa (Hill q = 2) and (d) Arthropod abundance. Significance levels are presented as: n.s = not significant, + = 0.10 > *p* > 0.05, * *p* <0.05, ** *p* <0.01, and *** *p* <0.001.

The effective number of common taxa (Hill q = 1) found within plots was not influenced by terpenoid diversity (Hill q = 0,1,2), average number of stems and average plant height (Fig. 4b, S7b & S8b & Table S4). Terpenoid abundance (*X*² = 4.7, p = 0.030) in plots however, reduced the effective number of common taxa found in plots (S9b & Table S4).

Average plant height (*X*² = 1.5, p = 0.213) did not significantly affect the effective number of dominant taxa observed within plots but the average number of stems (*X*² = 3.7, p = 0.056) had a marginal impact on the effective number of dominant taxa found in plots (Table S4).

None of the terpenoid diversity metrics (Hill q = 0,1,2) significantly influenced the effective number of dominant taxa found within plots (Fig. 4c, S7c, S8c & Table S4). Plot terpenoid abundance (*X*² = 6.2, p = 0.013), however, reduced the effective number of dominant taxa associated with plots (Fig. S9c & Table S4).

Average plant height increased arthropod abundance within plots, but terpenoid diversity (Hill q = 0,1,2), terpenoid abundance and average number of stems had no impact on arthropod abundance in plots (Fig. 4d, S7d, S8d & Table S4).

## Discussion

In this study, we investigated the influence of plot-level chemodiversity and its components (α, β and γ-chemodiversity) on aboveground arthropod communities. We found that independent of the mean terpenoid (α-) diversity of chemotypes present within plots, the presence of certain Tansy chemotypes and plot-level chemotype richness (β-chemodiversity) significantly impacted aboveground arthropod community assemblage. We also show that terpenoid abundance (i.e. the potential concentration of terpenoids produced by plants) has significant influence on arthropod community diversity within plots.

### α-Chemodiversity does not impact arthropod communities

We found that the presence of the Chrys-acet and Mixed-high chemotypes within plots reduced the effective number of common and dominant arthropod taxa associated with plots. Despite the clear impact of these two chemotypes on arthropod diversity, the effects seem not to be related to terpenoid α-chemodiversity. For instance, the Chrys-acet chemotype whose is profile dominated by chrysanthenyl acetate, has a relatively high terpenoid richness when compared with the other chemotypes within our study (Neuhaus-Harr et al. 2024), whilst the chemotype Mixed-high which contains more evenly distributed terpenoids has comparably low mean terpenoid richness, this suggests that the α-chemodiversity of plants present in a plot, *per se*, is an unlikely driver of aboveground arthropod communities at the plot level. Instead, the identity of chemotypes is essential, as certain arthropods are well adapted to and show a clear preference for particular chemotypes selecting them over others, as has been shown in other studies (Neuhaus Harr et al., 2024; Ojeda Prieto, Moreno, et al., 2025; Ziaja & Müller, 2023). For example, in field surveys of different Tansy plants, Bálint et al., (2016) observed clear chemotype preferences of two aphid predators, aphid specialist beetle, *Coccinella septempunctata,* was frequently observed on camphor-dominated chemotypes whilst polyphagous spider, *Pisaura mirabilis* was often found on borneol-dominated plants. Moreover, both chemotypes Chrys-acet and Mixed-high exhibit high terpenoid abundance implying that apart from the identity of the chemotype, the potential amount of terpenoids expressed by the plant also influences interacting arthropods. A similar trend was observed on *Senecio jacobaea* plants, where overall damage to plants by herbivores was positively associated with the concentration of pyrrolizidine alkaloids in leaves (Macel & Klinkhamer, 2010). Neuhaus Harr et al., (2024) also found that Tansy specialist aphid *Macrosiphoniella tanaceteria* showed a preference for high concentration chemotypes in choice assays.

### β-Chemodiversity impacts on arthropod communities vary by analytical approach

Contrary to our hypothesis on β-chemodiversity (H2), our studies revealed no clear relationship between dissimilarities in leaf terpenoid composition of plants within plots and the dissimilarities of arthropod groups found within plots. Average plot dissimilarity also had no impact on the diversity and abundance of arthropod groups found within plots, demonstrating that differences in leaf terpenoid composition of the individual plants within our plots did not drive arthropod community assemblage. While terpenoid emission in plots may differ from stored leaf terpenoids, overlaps exist in their composition, these overlapping compounds are probably constitutively emitted from stored terpenoids reserves and make up the plant chemotype which associated organisms’ encounter (Clancy et al., 2016; Ziaja & Müller, 2025). Our study comes with two caveats; firstly, the terpenoid profiles have been measured at the start of the experiment, and these values have been used as proxies. However, given the fact that repeated measurement of cuttings from the same genotypes over the years have revealed that these chemotypes are chemically consistent (Newrzella et al., 2026), this should not be a concern. Secondly, the terpenoid profiles reflect stored terpenoids, which may have different impacts on arthropod attraction and repellence than emitted volatile terpenoids. This is an area where future work measuring both leaf terpenoid and headspace volatile terpenoid emission at the time of insect collection could help unravel their respective roles in driving arthropod assemblage.

Interestingly, determining β-chemodiversity with plot-level chemotype richness revealed a negative relationship between chemotype richness and the effective number of dominant arthropod taxa found in plots, indicating that having more diverse neighbours selects for specific arthropod taxa rather increasing overall taxa numbers. This is supported by findings by Richards et al., (2015), who found that increased phytochemical diversity in *Piper* plants promotes specialized herbivore species, and that chemical diversity in plants is a direct result of herbivore pressure by these specialized arthropods. In our study, we observed that Nematocera, a taxonomic group predominantly composed of Tansy-specialized flower midges (Cecidomyiidae), increased as chemotype richness increased. These midges may have co-evolved with Tansy plants and hence may select for plots with more variable emissions as these plots would have fewer competing taxa and a higher diversity of food resources. Other canopy arthropod taxa which may be more transient generalists may not be well adapted to these conditions, preferring specific plots with more uniform profiles. Many of the observed taxa are commensalists, meaning that they do not directly harm or benefit the plant, but may benefit from living in the plant canopy in form of habitat, refuge, or the provision of non-plant resources.

### Chemical abundance at plot-level instead of γ-chemodiversity shapes arthropod communities

Notably, we found no effects of plot terpenoid diversity on arthropod community diversity and abundance. This is contrary to the few other studies that report effects of community-level chemodiversity on arthropod communities, particularly on herbivore dynamics (Fernandez Conradi et al., 2022; Calixto et al., 2025; Salazar et al., 2016; Wang et al., 2023). For example, lower herbivore damage has been observed on plant communities with high pooled chemodiversity at lower elevations, a pattern that reverses at higher elevations (Fernandez Conradi et al., 2022). We speculate that certain stored terpenoids may be inactive and do not effectively contribute to the plant chemotype perceived by associated arthropods (Iason et al., 2011; Wetzel & Whitehead, 2020), as such arthropod communities may be responding to select subsets of terpenoids or terpenoid blends and not the pooled cumulative terpenoid diversity within plots. Further studies are required to fully understand which particular terpenoids or terpenoid blends are actively shaping associated arthropod communities at specific timepoints.

We observed a negative relationship between plot terpenoid abundance and the effective number of common and dominant taxa found in plots. Terpenoid abundance, which is a proxy for relative terpenoid concentration produced per unit leaf biomass, has been reported by several studies to be used as a cue for host finding by certain arthropods (Calixto et al., 2025; Neuhaus Harr et al., 2024; Setordjie et al., 2026). According to Calixto et al., (2025), certain leaf gallers have been found to have a positive relationship with terpenoid concentration in *Monandra fistucola* plants. In our study, we found that increased terpenoid abundance led to a reduction in most arthropod taxa except for Nematocera, which were dominated by specialized flower midges that may use terpenoid concentration as a host-finding cue, as has been found in other plant systems (Badenes-Pérez et al., 2020; Poelman et al., 2009). We theorize that the use of terpenoid abundance as a host finding cue is highly taxon-specific and is especially used by specialist taxa. Generalist arthropod taxa are likely to avoid plots with high terpenoid abundance as they may not be well adapted to such conditions. Nevertheless, plant terpenoid composition can vary due to environmental conditions, future work should explore the role of terpenoid abundance in shaping arthropod communities at different time points and plant developmental stages.

### Other factors influencing arthropod community assemblage

Even though suction sampling is a wildly used and effective technique for determining arthropod diversity and abundance, its efficiency varies largely with arthropod taxa and form as well as vegetation type (Sanders & Entling, 2011; Doxon et al., 2011). Less than 30% efficiency has been reported for ground dwelling arthropods like Collembola, Isopoda, Diplopoda, and Formicidae (Sanders & Entling, 2011). Arthropods like Coleoptera and Staphylinidae capable of hiding under litter and soil may also be missed during sampling, as well as larger and highly mobile arthropods (Sanders & Entling, 2011; Doxon et al., 2011). Aphids, which happen to be the most abundant herbivores on Tansy, are also reported to be under sampled when using suction due to their ability to attach closely to plant stems using their claws and other adhesive organs (Lees & Hardie, 1988; Dixon et al., 1990). This however does not negate our conclusions as we clearly observe effects Tansy chemodiversity on Nematocera, a taxon composed of specialized tansy flower midges, highlighting a close association and adaptation to Tansy, which is absent for all other taxa regardless of their abundance.

## Conclusion

Intraspecific chemodiversity in plant communities play a crucial role in shaping associated arthropod communities, how its spatial components (α, β and γ-chemodiversity) contribute to this role is complex and dependent on interacting taxa. Our study reveals that rather than terpenoid diversity and leaf terpenoid dissimilarities, chemotype identity, plot-level chemotype richness and terpenoid abundance are the factors influencing arthropod communities in plant canopies by selecting for well adapted and specialist arthropods. Future studies should however delve into mechanisms behind the evolution of distinct chemotypes, their distribution in nature and their importance in ecological relationships.

## Acknowledgements

We express our gratitude to the Jena Experiment management, particularly Anne Ebeling and Nico Eisenhauer for allowing us to maintain our experiment on-site, and to the local gardening team for lending their equipment and assisting with information where necessary. We are also thankful for the support of Walter D. Espindola and Maximilian Bröcher for helping with preliminary instructions and equipment set-up and to Rafael Achury and Petra Freynhagen for assisting with sorting instructions.

## Author contributions

W.W. Weisser designed the experiment. R. Heinen propagated the plants and R. Heinen, L. Ojeda-Prieto and W. W. Weisser set-up the experimental field. E. A. Setordjie and L. Ojeda-Prieto maintained the field experiment. E. A. Setordjie, A. Neuhaus and S. Chandra sampled arthropods. E. A. Setordjie, S. Chandra and L. Ojeda-Prieto sorted arthropod samples into (sub)orders. E. A. Setordjie organized and analysed data. E. A. Setordjie wrote the manuscript with the guidance of R. Heinen. L. Ojeda-Prieto, A. Neuhaus, S. Chandra, and W.W. Weisser provided critical feedback on earlier drafts. All authors contributed to the final version and approved the final submission.

