## Supplementary materials for "Unravelling the role of plant alpha-, beta-, and gamma- chemodiversity in shaping canopy insects of plants within the same species"

ORCIDs:

Elikplim Aku Setordjie: <https://orcid.org/0000-0002-8516-7736>

Annika Neuhaus: <https://orcid.org/0009-0009-6093-4108>

Sunayana Chandra: <https://orcid.org/0009-0001-6840-7187>

Lina Ojeda-Prieto: <https://orcid.org/0009-0003-6069-901X>

Wolfgang W. Weisser: <https://orcid.org/0000-0002-2757-8959>

Robin Heinen: <https://orcid.org/0000-0001-9852-1020>

**Supplementary tables:**

**Table S1**: Model output for the effect of chemotype presence

Degrees of freedom (df), Wald’s Chi-square statistics and p-values are reported in the table**.**

Arthropod taxon richness (Hill q = 0), Effective number of common taxa (Hill q = 1), Effective number of dominant taxa (Hill q = 2).

|  |  | Arthropod | | | |
| --- | --- | --- | --- | --- | --- |
|  |  | Hill q = 0 | Hill q = 1 | Hill q = 2 | Abundance |
|  | d.f. | X^2^ (p-value) | X^2^ (p-value) | X^2^ (p-value) | X^2^ (p-value) |
| Average number of stems | 1 | 1.0 (0.329) | 0.2 (0.685) | 0.3 (0.605) | 0.0 (0.935) |
| Average plant height | 1 | 0.2 (0.673) | 1.6 (0.210) | 2.3 (0.128) | 5.3 (0.021) |
| Athu-Bthu | 1 | 0.7 (0.402) | 0.0 (0.893) | 0.1 (0.725) | 0.1 (0.721) |
| Average number of stems | 1 | 0.9 (0.351) | 0.2 (0.655) | 0.2 (0.673) | 0.0 (0.885) |
| Average plant height | 1 | 0.1 (0.823) | 1.0 (0.316) | 2.4 (0.121) | 4.1 (0.043) |
| Bthu-low | 1 | 0.0 (0.831) | 0.4 (0.538) | 0.2 (0.659) | 0.2 (0.664) |
| Average number of stems | 1 | 1.0 (0.324) | 0.2 (0.631) | 0.3 (0.579) | 0.0 (0.907) |
| Average plant height | 1 | 0.1 (0.804) | 0.9 (0.341) | 1.4 (0.238) | 4.0 (0.046) |
| Bthu-high | 1 | 0.0 (0.899) | 1.0 (0.316) | 1.6 (0.210) | 1.1 (0.294) |
| Average number of stems | 1 | 0.8 (0.360) | 0.3 (0.562) | 0.5 (0.477) | 0.0 (0.884) |
| Average plant height | 1 | 0.1 (0.770) | 1.6 (0.207) | 2.3 (0.126) | 5.2 (0.023) |
| Chrys-acet | 1 | 0.8 (0.360) | 4.2 (0.041) | 7.0 (0.008) | 0.6 (0.428) |
| Average number of stems | 1 | 0.6 (0.430) | 0.0 (0.848) | 0.1 (0.715) | 0.0 (0.874) |
| Average plant height | 1 | 0.1 (0.800) | 1.6 (0.201 | 2.2 (0.138) | 4.8 (0.028) |
| Mixed-low | 1 | 0.0 (0.832) | 0.1 (0.773) | 0.0 (0.900) | 0.0 (0.856) |
| Average number of stems | 1 | 0.8 (0.372) | 1.2 (0.266) | 1.9 (0.167) | 0.4 (0.515) |
| Average plant height | 1 | 0.0 (0.829) | 0.4 (0.548) | 0.6 (0.443) | 2.8 (0.094) |
| Mixed-high | 1 | 0.0 (0.862) | 3.6 (0.056) | 4.8 (0.029) | 2.4 (0.124) |

**Table S2**: Model output for the effect of average plot dissimilarity

Degrees of freedom (df), Wald’s Chi-square statistics and p-values are reported in the table**.**

Arthropod taxon richness (Hill q = 0), Effective number of common taxa (Hill q = 1), Effective number of dominant taxa (Hill q = 2).

|  |  | Arthropod | | | |
| --- | --- | --- | --- | --- | --- |
|  |  | Hill q = 0 | Hill q = 1 | Hill q = 2 | Abundance |
|  | d.f. | X^2^ (p-value) | X^2^ (p-value) | X^2^ (p-value) | X^2^ (p-value) |
| Average number of stems | 1 | 0.9 (0.344) | 0.1 (0.712) | 0.2 (0.663) | 0.0 (0.978) |
| Average plant height | 1 | 0.1 (0.776) | 1.5 (0.214) | 2.2 (0.140) | 5.2 (0.023) |
| Average plot dissimilarity | 1 | 0.3 (0.558) | 0.1 (0.795) | 0.2 (0.643) | 0.7 (0.391) |

**Table S3**: Model output for the effect of plot-level chemotype richness.

Degrees of freedom (df), Wald’s Chi-square statistics and p-values are reported in the table**.**

Arthropod taxon richness (Hill q = 0), Effective number of common taxa (Hill q = 1), Effective number of dominant taxa (Hill q = 2).

|  |  | Arthropod | | | |
| --- | --- | --- | --- | --- | --- |
|  |  | Hill q = 0 | Hill q = 1 | Hill q = 2 | Abundance |
|  | d.f. | X^2^ (p-value) | X^2^ (p-value) | X^2^ (p-value) | X^2^ (p-value) |
| Average number of stems | 1 | 0.9 (0.342) | 0.2 (0.631) | 0.3 (0.556) | 0.0 (0.885) |
| Average plant height | 1 | 0.0 (0.869) | 0.9 (0.332) | 1.3 (0.252) | 4.1 (0.043) |
| Plot-level chemotype richness | 1 | 0.5 (0.486) | 2.4 (0.120) | 4.0 (0.045) | 1.7 (0.192) |

**Table S4**: Model output for the effect of plot terpenoid diversity metrics

Degrees of freedom (df), Wald’s Chi-square statistics and p-values are reported in the table**.**

Arthropod taxon richness (Hill q = 0), Effective number of common taxa (Hill q = 1), Effective number of dominant taxa (Hill q = 2).

|  |  | Arthropod | | | |
| --- | --- | --- | --- | --- | --- |
|  |  | Hill q = 0 | Hill q = 1 | Hill q = 2 | Abundance |
|  | d.f. | X^2^ (p-value) | X^2^ (p-value) | X^2^ (p-value) | X^2^ (p-value) |
| Average number of stems | 1 | 1.0 (0.319) | 0.1 (0.710) | 0.2 (0.656) | 0.0 (0.984) |
| Average plant height | 1 | 0.1 (0.792) | 1.4 (0.229) | 2.0 (0.159) | 4.6 (0.032) |
| Plot terpenoid richness  (Hill q = 0) | 1 | 0.0 (0.915) | 0.0 (0.849) | 0.1 (0.701) | 0.5 (0.496) |
| Average number of stems | 1 | 1.0 (0.320) | 0.2 (0.694) | 0.2 (0.619) | 0.0 (0.945) |
| Average plant height | 1 | 0.1 (0.810) | 1.1 (0.289) | 1.6 (0.201) | 4.3 (0.038) |
| Effective number of common terpenoids in plot  (Hill q = 1) | 1 | 0.1 (0.797) | 1.5 (0.222) | 1.8 (0.182) | 1.7 (0.197) |
| Average number of stems | 1 | 1.0 (0.323) | 0.2 (0.658) | 0.3 (0.579) | 0.0 (0.908) |
| Average plant height | 1 | 0.1 (0.791) | 1.0 (0.307) | 1.6 (0.207) | 4.3 (0.039) |
| Effective number of dominant terpenoids in plot (Hill q = 2) | 1 | 0.0 (0.918) | 1.8 (0.183) | 1.8 (0.182) | 1.3 (0.246) |
| Average number of stems | 1 | 0.5 (0.461) | 2.4 (0.121) | 3.7 (0.056) | 0.3 (0.613) |
| Average plant height | 1 | 0.1 (0.801) | 0.9 (0.334) | 1.5 (0.213) | 4.5 (0.034) |
| Plot terpenoid abundance | 1 | 0.1 (0.753) | 4.7 (0.030) | 6.2 (0.013) | 0.6 (0.434) |

**Supplementary figures:**

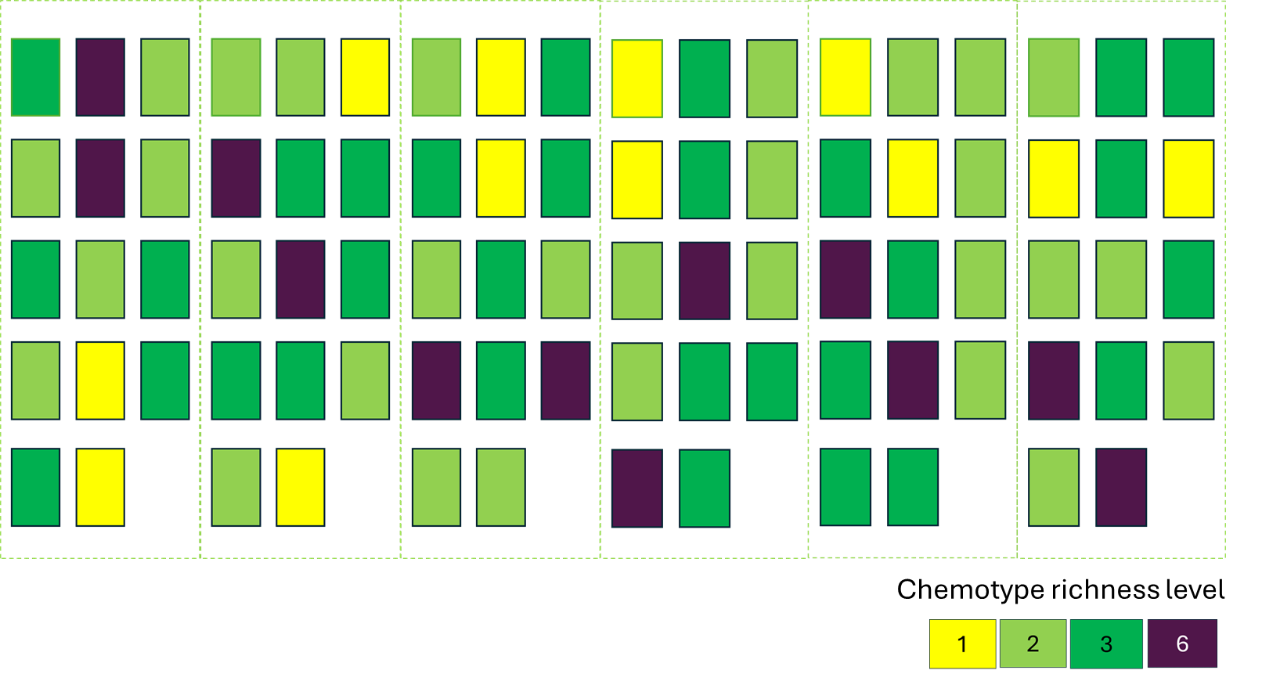

**Figure S1:** Graphical representation of field layout in Jena, Germany, consisting of 84 plots (14 plots per block, blocks demarcated by dashed lines). Each rectangle represents a plot.

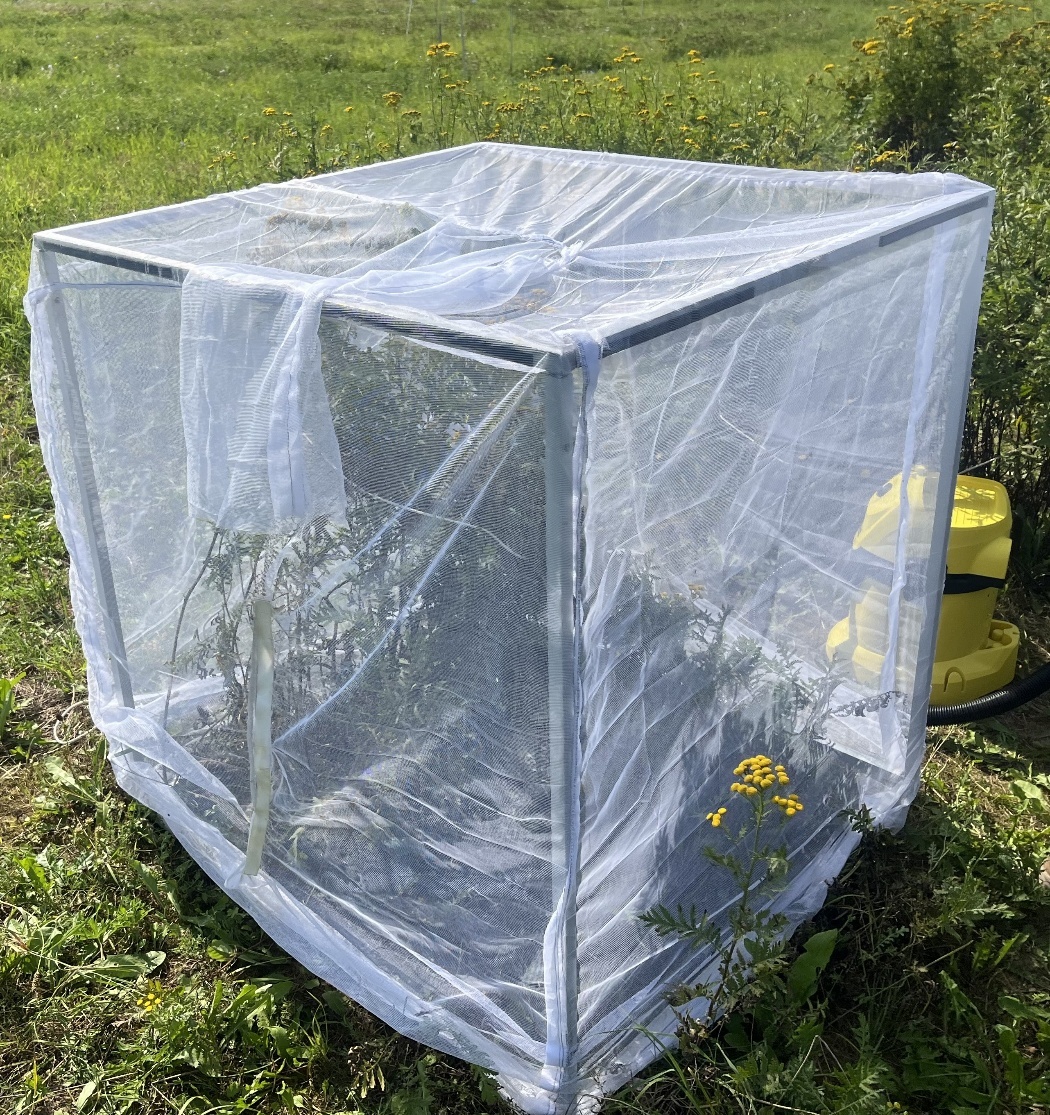

**Figure S2**: Picture of netted cage (placed on a plot in field) and vacuum (yellow device) used for sampling.

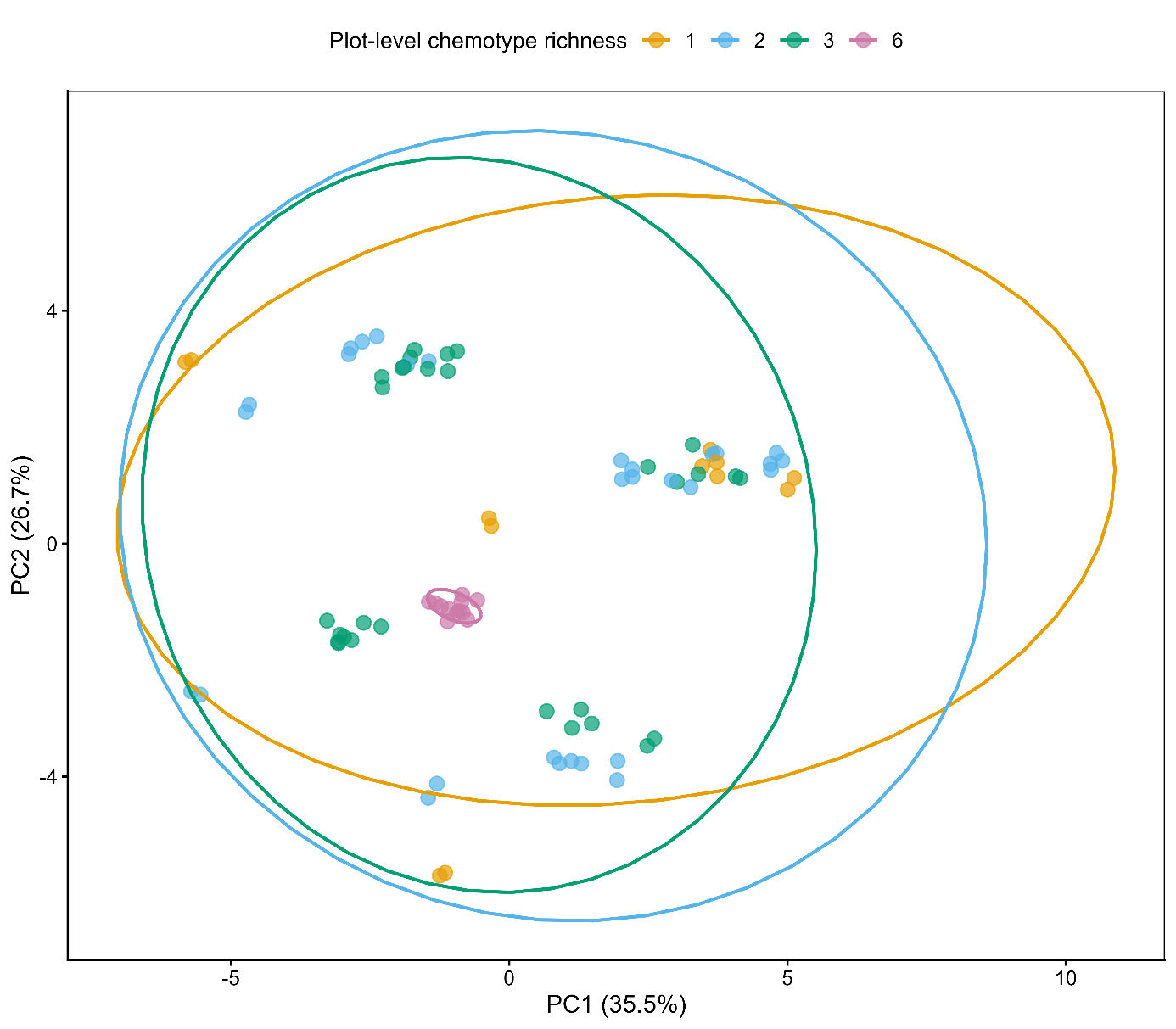

**Figure S3:** Principal component analysis showing terpenoid variations across different plot chemotype richness levels (NB: Terpenoid abundances were log+1 transformed prior to principal component analysis).

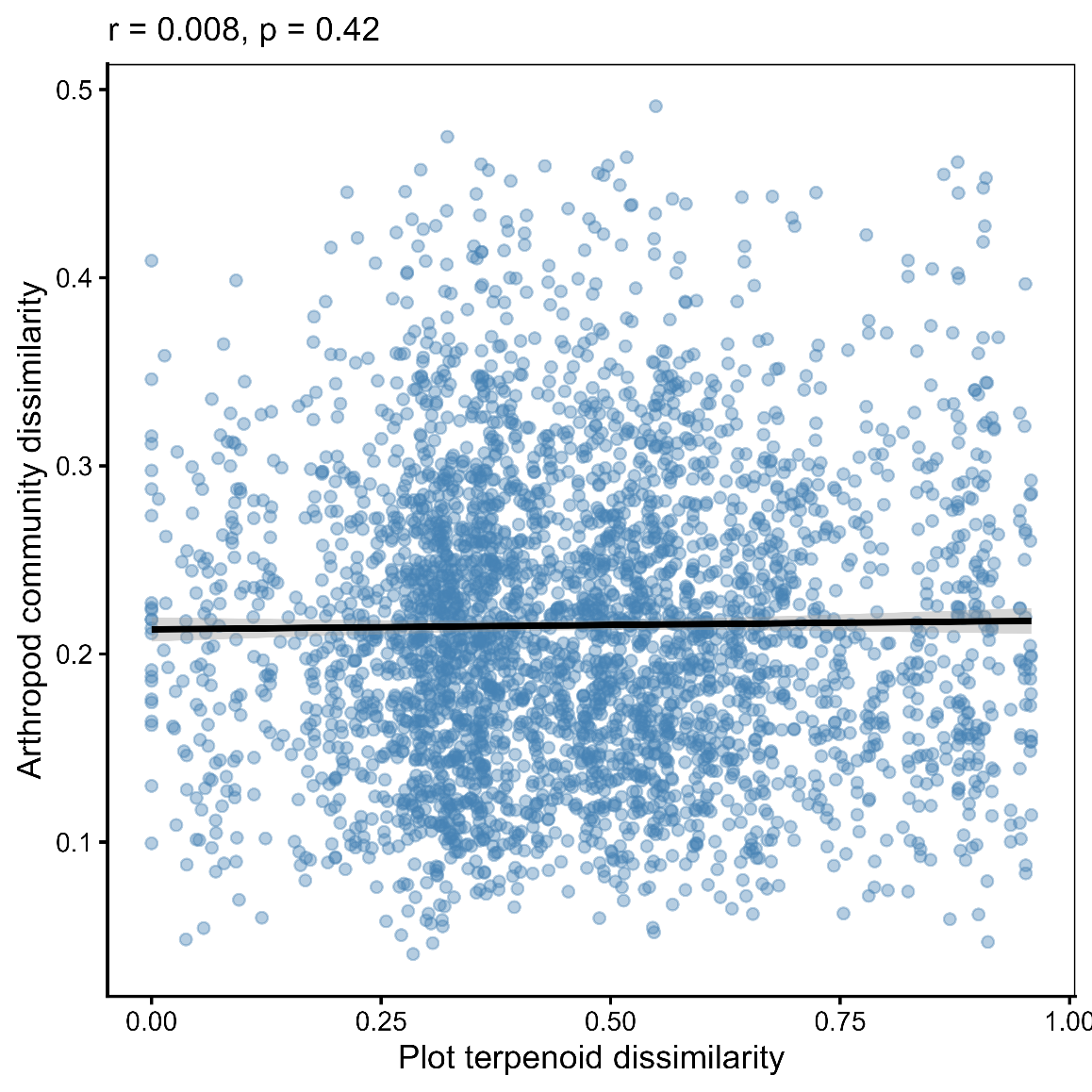

**Figure S4:** Correlation between plot terpenoid dissimilarity and arthropod community dissimilarity after Mantel test ran at 9999 permutations.

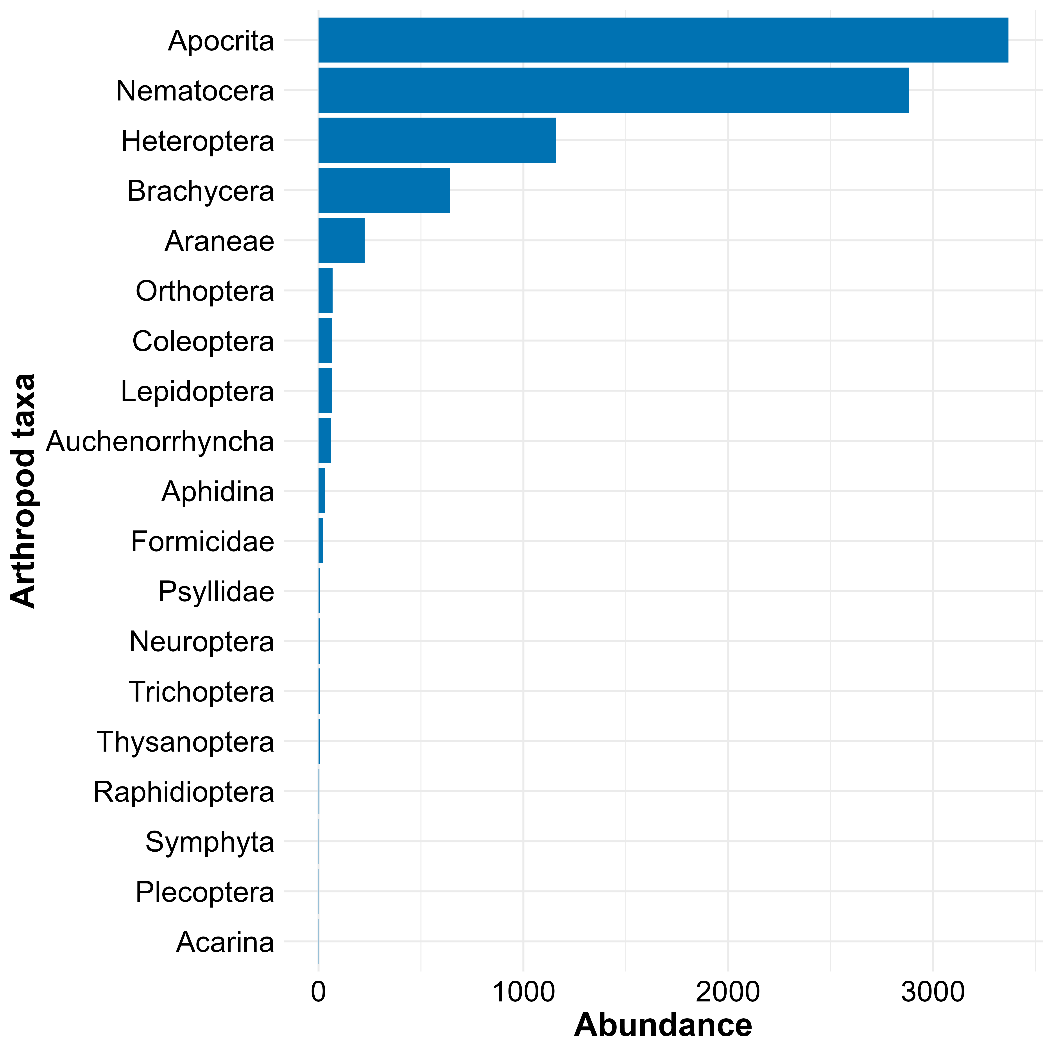

**Figure S5**: Abundance of different arthropod taxa within our samples

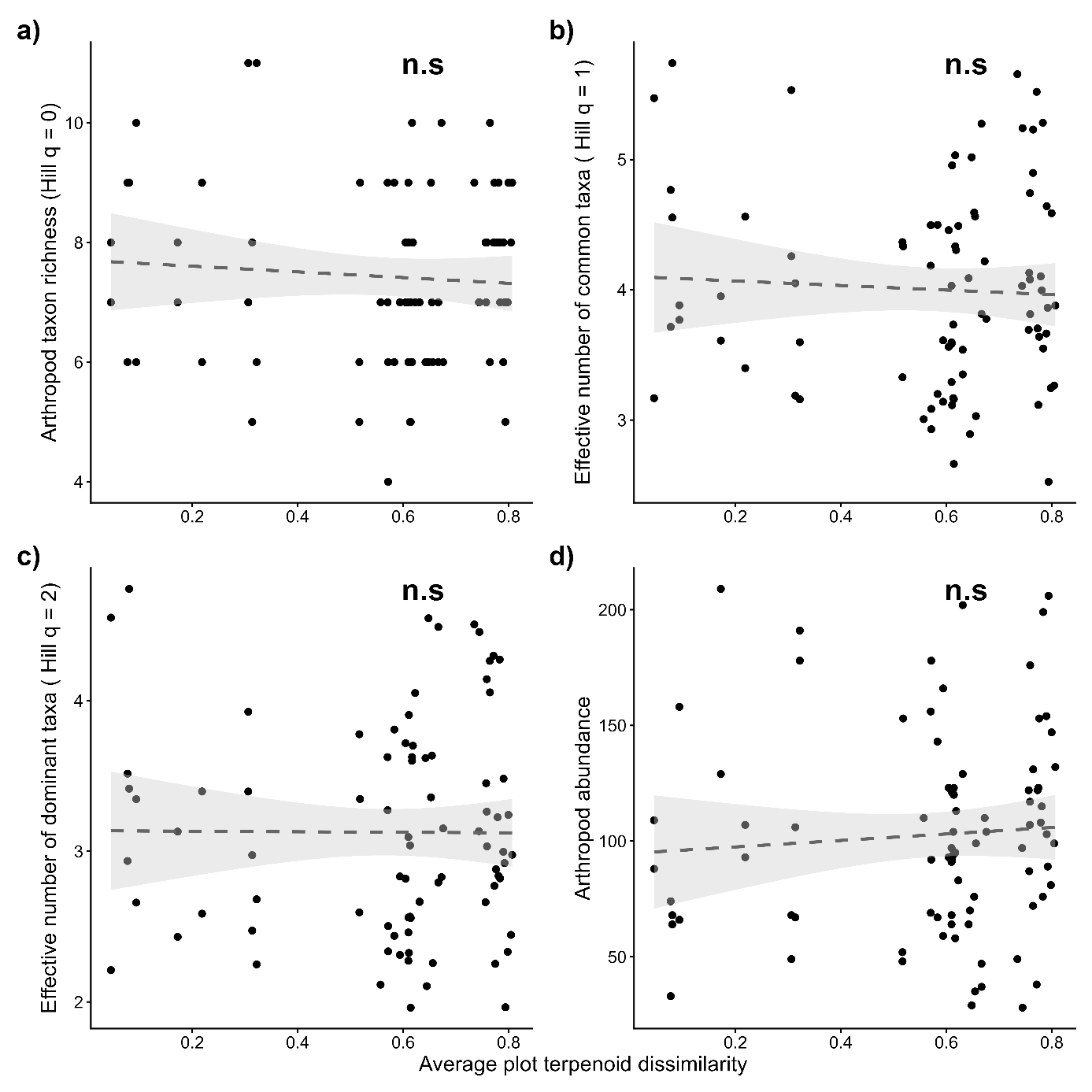

**Figure S6:** Effect of average plot terpenoid dissimilarity on (a) Arthropod taxon richness (Hill q = 0), (b) Effective number of common taxa ( Hill q = 1), (c) Effective number of dominant taxa (Hill q = 2) and (d) Arthropod abundance. Significance levels are presented as: n.s = not significant, + = 0.10 > *p >* 0.05, * *p* <0.05, ** *p* <0.01, and *** *p* <0.001.

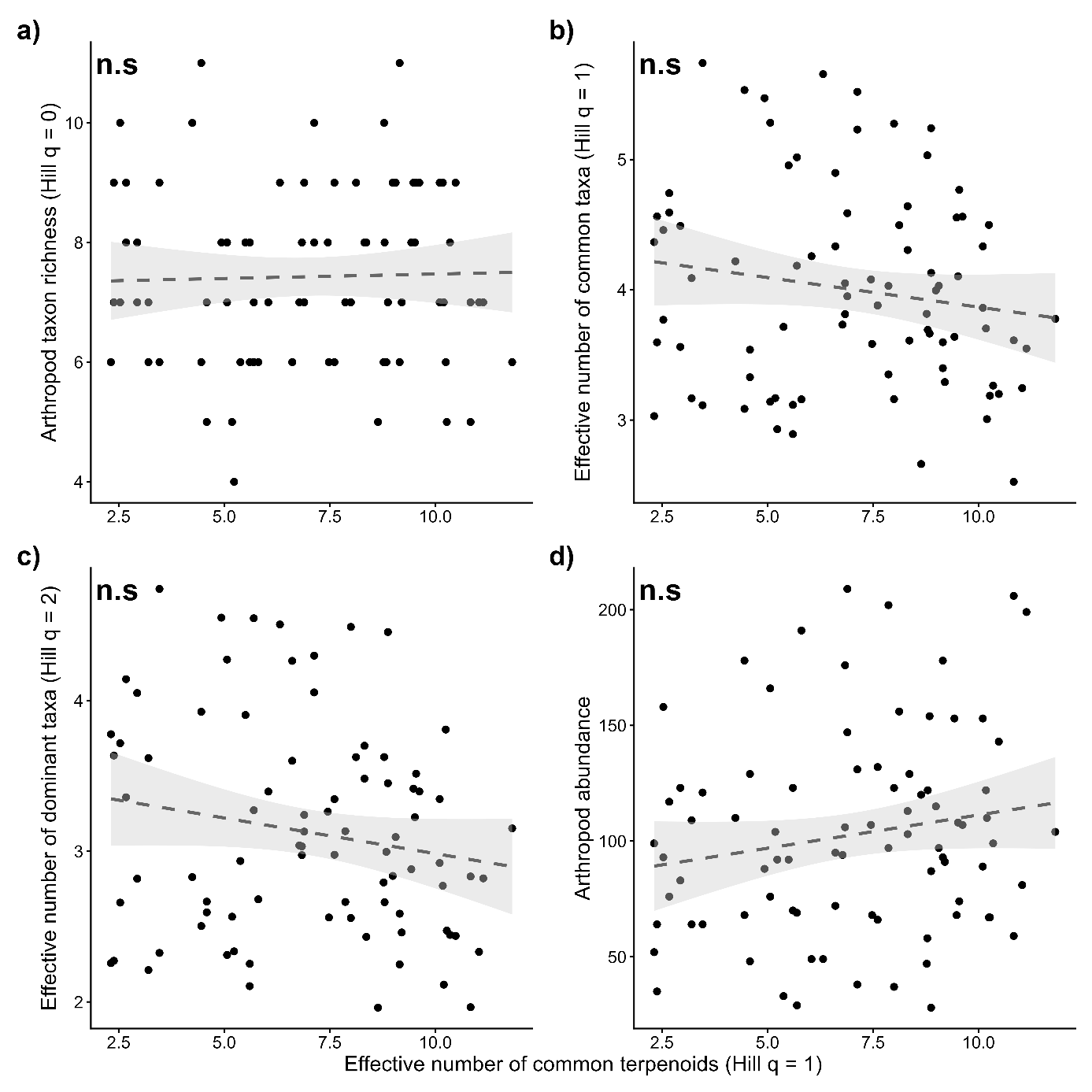

**Figure S7:** Effect of the effective number of common terpenoids within plots (Hill q = 1) on (a) Arthropod taxon richness (Hill q = 0), (b) Effective number of common taxa (Hill q = 1), (c) Effective number of dominant taxa (Hill q = 2) and (d) Arthropod abundance. Significance levels are presented as: n.s = not significant, + = 0.10 > *p* > 0.05, * *p* <0.05, ** *p* <0.01, and *** *p* <0.001.

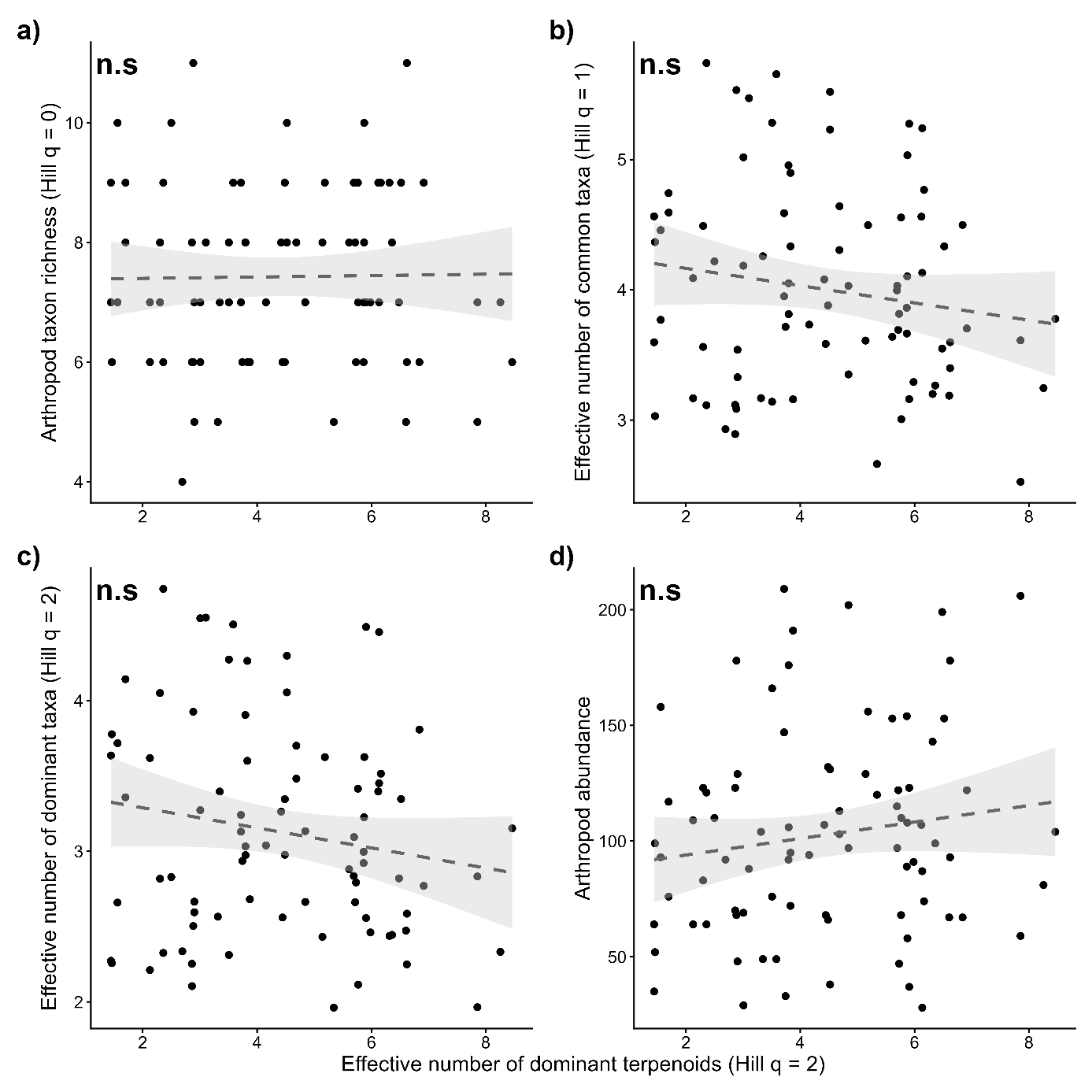
**Figure S8:** Effect of the effective number of dominant terpenoids within plots (q = 2) on (a) Arthropod taxon richness (q = 0), (b) Effective number of common taxa (q = 1), (c) Effective number of dominant taxa (q = 2) and (d) Arthropod abundance. Significance levels are presented as: n.s = not significant, + = 0.10 > *p* > 0.05, * *p* <0.05, ** *p* <0.01, and *** *p* <0.001.

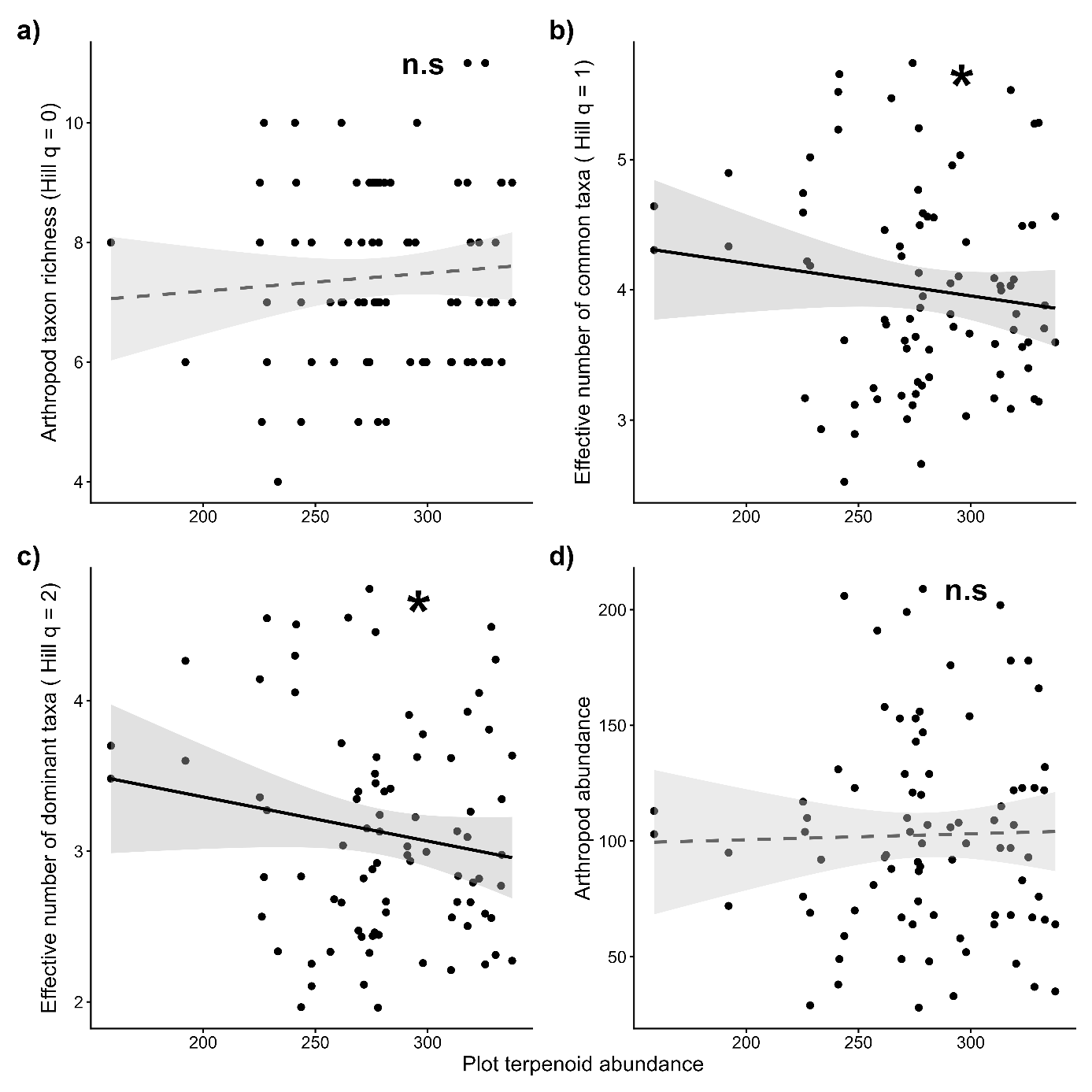

**Figure S9**: Effect of plot terpenoid abundance on (a) Arthropod taxon richness (Hill q = 0), (b) Effective number of common taxa (Hill q = 1), (c) Effective number of dominant taxa (Hill q = 2) and (d) Arthropod abundance. Significance levels are presented as: n.s = not significant, + = 0.10 > *p* > 0.05, * *p* <0.05, ** *p* <0.01, and *** *p* <0.001.
